# The leucine-rich repeat receptor kinase QSK1 is a novel regulator of PRR-RBOHD complex and is employed by the bacterial effector HopF2*_Pto_* to modulate plant immunity

**DOI:** 10.1101/2023.12.13.571443

**Authors:** Yukihisa Goto, Yasuhiro Kadota, Malick Mbengue, Jennifer D Lewis, Hidenori Matsui, Noriko Maki, Jan Sklenar, Paul Derbyshire, Arisa Shibata, Yasunori Ichihashi, David S. Guttman, Hirofumi Nakagami, Takamasa Suzuki, Frank L.H. Menke, Silke Robatzek, Darrell Desveaux, Cyril Zipfel, Ken Shirasu

## Abstract

Plants detect pathogens using cell-surface pattern recognition receptors (PRRs) like EFR and FLS2, which recognize bacterial EF-Tu and flagellin, respectively. These PRRs, belonging to the leucine-rich repeat receptor kinase (LRR-RK) family, activate the production of reactive oxygen species via the NADPH oxidase RBOHD. The PRR-RBOHD complex is tightly regulated to prevent unwarranted or exaggerated immune responses. However, certain pathogenic effectors can subvert these regulatory mechanisms, thereby suppressing plant immunity. To elucidate the intricate dynamics of the PRR-RBOHD complex, we conducted a comparative co-immunoprecipitation analysis using EFR, FLS2, and RBOHD. We identified QSK1, an LRR-RK, as a novel component of the PRR-RBOHD complex. QSK1 functions as a negative regulator of PRR-triggered immunity (PTI) by downregulating the abundance of FLS2 and EFR. QSK1 is targeted by the bacterial effector HopF2*_Pto_*, a mono-ADP ribosyltransferase, resulting in the reduction of FLS2 and EFR levels through both transcriptional and transcription-independent pathways, thereby inhibiting PTI. Furthermore, HopF2*_Pto_* reduces transcript levels of *PROSCOOP* genes encoding important stress-regulated phytocytokines and their receptor MIK2. Importantly, HopF2*_Pto_* requires QSK1 for its accumulation and virulence functions within plants. In summary, our results provide novel insights into the mechanism by which HopF2*_Pto_* employs QSK1 to desensitize plants to pathogen attack.

**One Sentence Summary:** QSK1, a novel component in the plant immune receptor complex, downregulates these receptors and phytocytokines, and is exploited by bacterial effector HopF2*_Pto_* to desensitize plants to pathogen attack.

## Introduction

Plants and pathogens are in a perpetual evolutionary arms race. A fundamental aspect of the plant’s defense mechanism lies in its capability to detect microbial molecules, particularly pathogen-associated molecular patterns (PAMPs) as well as endogenous danger molecules that are released from damaged or dying cells, known as damage-associated molecular patterns (DAMPs). These PAMPs and DAMPs are recognized by specialized cell-surface receptors known as pattern recognition receptors (PRRs) (Macho and Zipfel, 2014). Among those, leucine-rich repeat receptor-kinases (LRR-RKs) play a central role in the recognition of PAMPs and DAMPs. For instance, ELONGATION Factor-TU (EF-Tu) RECEPTOR (EFR) and FLAGELLIN SENSING2 (FLS2) detect bacterial EF-Tu and flagellin, respectively. The binding of flg22 or elf18 (the immunogenic peptides of flagellin or EF-Tu, respectively) to FLS2 and EFR induces their instant association with the coreceptor LRR-RK BRI1-ASSOCIATED RECEPTOR KINASE 1 (BAK1) and concomitant phosphorylation of both proteins to initiate PRR-triggered immunity (PTI) (Chinchilla et al., 2007; Heese et al., 2007; Roux et al., 2011). Subsequently, the PRR-BAK1 complex activates receptor-like cytoplasmic kinases (RLCKs) such as BOTRYTIS INDUCED KINASE 1 (BIK1) by phosphorylation (Lu et al., 2010; Zhang et al., 2010; Liu et al., 2013). PRRs further form a complex with the NADPH oxidase RESPIRATORY BURST OXIDASE HOMOLOG D (RBOHD), which is phosphorylated by activated BIK1, resulting in the rapid generation of reactive oxygen species (ROS) (Kadota et al., 2014; Li et al., 2014; Kadota et al., 2015). In addition, phosphorylated BIK1 activates Ca^2+^ channels, including OSCA1.3 (HYPEROSMOLALITY-GATED Ca^2+^-PERMEABLE CHANNEL1.3), CNGC2 (CYCLIC NUCLEOTIDE-GATED CHANNEL2), and CNGC4, particularly under specific Ca^2+^ concentrations (Tian et al., 2019; Thor et al., 2020). This activation leads to an increase in cytoplasmic Ca^2+^ concentration, subsequently stimulating Ca^2+^-dependent protein kinases (Boudsocq et al., 2010). Furthermore, BIK1 phosphorylates the non-canonical Gα protein, XLG2, facilitating its translocation to the nucleus. This phenomenon inhibits MUT9-like kinases, thereby removing the negative regulation of PTI (Liang et al., 2016; Ma et al., 2022). The remarkable orchestration of signal transduction within PRR complexes allows plants to mount swift and effective immune responses at the very site of infection.

To overcome effective plant immunity, the pathogens deploy virulence effectors to target and dampen immune signaling components (Dou and Zhou, 2012). Effectors with high immunomodulatory activities, especially those that suppress early PTI responses such as ROS production, MAPK activation, and Ca^2+^ influx, often target PRRs or their associated components. For example, AvrPto, a type III effector from *P. syringae*, directly inhibits the kinase activity of FLS2 and EFR (Xiang et al., 2008). AvrPtoB functions as an E3 ligase, catalyzing the polyubiquitination and degradation of FLS2, BAK1, and CERK1 (Goehre et al., 2008; Gimenez-Ibanez et al., 2009; Cheng et al., 2011). HopB1 associates with FLS2 and serves as a protease, cleaving activated BAK1 (Li et al., 2016). The *Xanthomonas campestris* effector AvrAC employs a unique uridylyl-transferase activity to impede the activation of BIK1 (Feng et al., 2012). These findings highlight the utility of effectors that suppress early PTI responses as valuable tools for identifying and confirming PRR complex components. Indeed, key regulators in the PRR complex, such as BIK1, and PBLs, were originally identified as targets of the bacterial effector AvrPphB, which possesses cysteine protease activity (Zhang et al., 2010). A comprehensive investigation of PRR complex components in conjunction with virulence effectors will shed light on the essential regulatory mechanisms governing PRR complexes and uncover how pathogens manipulate the PRR complex to enhance their virulence.

In this study, we used comparative immunoprecipitation (IP) analysis of EFR, FLS2, and RBOHD followed by mass-spectrometry (IP-MS) to identify components of mature PRR-RBOHD complexes situated at the plasma membrane. This investigation led to the identification of QIAN SHOU KINASE1 (QSK1), an LRR-RK, as a new component of this complex. Intriguingly, QSK1 plays a negative regulatory role in PTI, possibly by controlling the steady-state levels of PRRs. Our interaction assays further revealed an association between the bacterial effector HopF2*_Pto_* and QSK1. HopF2*_Pto_*, a mono-ADP ribosyltransferase, reduces PRR protein levels through both transcriptional and transcription-independent mechanisms. Moreover, HopF2*_Pto_* disrupts the signaling induced by SERINE RICH ENDOGENOUS PEPTIDE (SCOOP) phytocytokines. Importantly, the accumulation and virulence activities of HopF2*_Pto_*within plants rely on QSK1. In summary, our findings provide insights into the mechanisms by which QSK1 modulates PRR abundance and how HopF2*_Pto_* exploits QSK1 to render plant cells insensitive to PAMPs, DAMPs and SCOOP phytocytokines.

## Results

### Identification of QSK1, a novel component of PRR-RBOHD complexes

To isolate components specific to mature PRR-RBOHD complexes at the plasma membrane, we employed a comparative IP-MS strategy with EFR, FLS2, and RBOHD. Given the distinct protein structures of PRRs and RBOHD, it is likely that associated regulatory proteins involved in protein modification, maturation, transport, and degradation processes differ. Therefore, proteins that can associate with EFR, FLS2, and RBOHD are the most likely candidates to be associated with mature PRR-RBOHD complexes. To mitigate potential false positives resulting from sticky proteins, we implemented two different IP systems: magnetic and agarose beads. Through IP of FLS2-GFP from the Arabidopsis *pFLS2:FLS2-GFP* line using anti-GFP magnetic beads, we identified 118 FLS2-associated candidates (Supplemental Data Set S1_1). We had previously performed an IP of EFR-GFP using anti-GFP magnetic beads from the *efr-1/pEFR:EFR-GFP* line, identifying 42 candidate EFR-associated proteins (Supplemental Data Set S1_2) (Kadota et al., 2014). Moreover, we previously identified 451 candidate RBOHD-associated proteins through IP of 3xFLAG-RBOHD from the *rbohD*/*pRBOHD:3xFLAG-RBOHD* line by using Anti-FLAG agarose and eluted 3xFLAG-RBOHD with free 3xFLAG peptides (Supplemental Data Set S1_3) (Goto et al., 2023). Venn diagram analysis of these candidates pinpointed thirteen proteins commonly associated with FLS2, EFR, and RBOHD (Fig. 1), including known components of PRR complexes such as BAK1 (Chinchilla et al., 2007; Heese et al., 2007; Roux et al., 2011), IMPAIRED OOMYCETE SUSCEPTIBILITY 1 (IOS1) (Yeh et al., 2016), AUTOINHIBITED Ca^2+^-ATPASE 10 (ACA10) (Frei dit Frey et al., 2012), and RBOHD (Kadota et al., 2014; Li et al., 2014). Additionally, several proteins are known to accumulate in detergent-resistant membrane compartments in response to flg22, including QSK1, ACA10, SYNTAXIN OF PLANTS 71 (SYP71), HYPERSENSITIVE INDUCED REACTION1 (HIR1), HIR4, and REMORIN 1.2 (REM1.2) (Keinath et al., 2010). These results validate the effectiveness of our comparative IP-MS approach for identifying members of mature PRR-RBOHD complexes.

**Figure 1.**
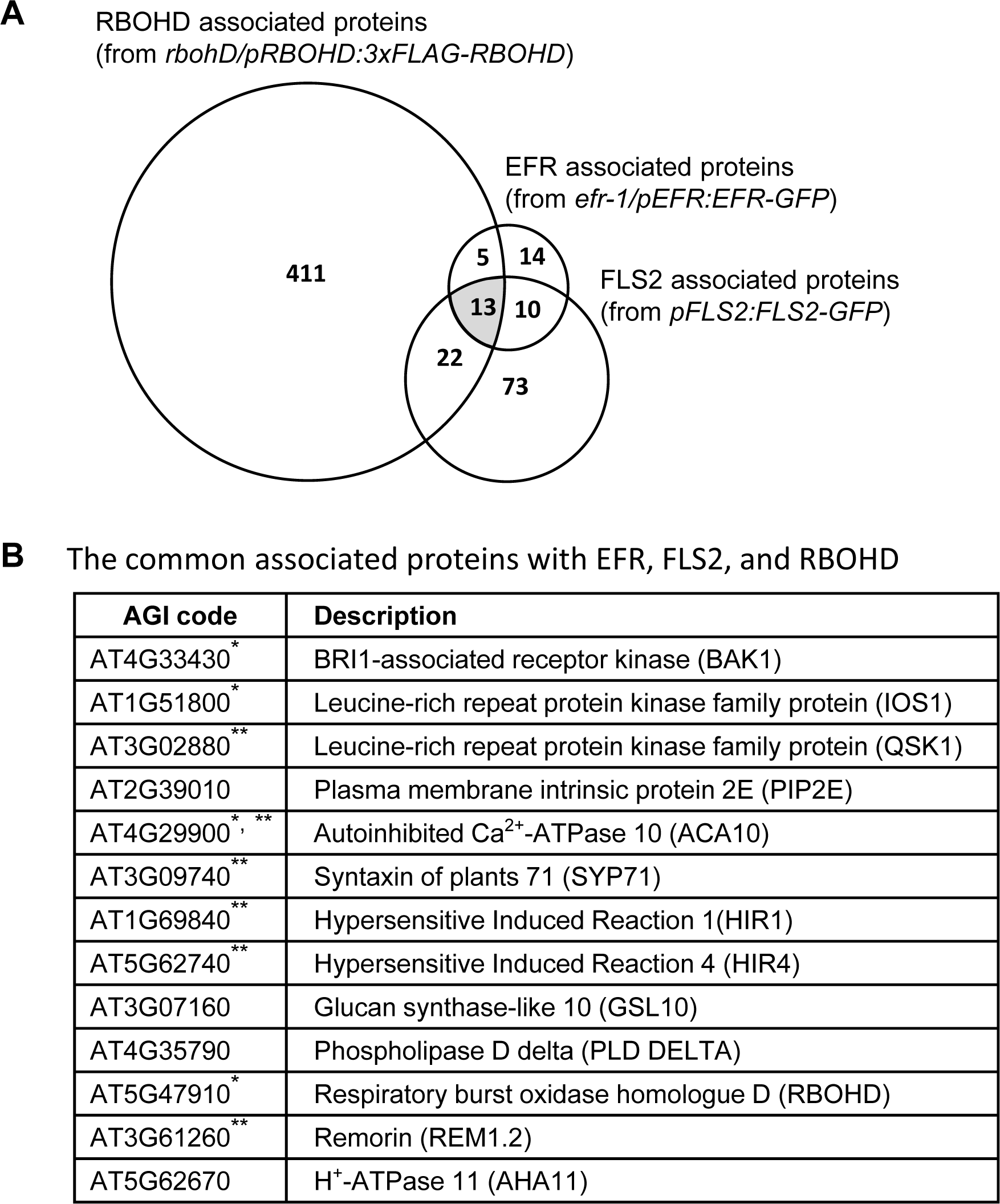
Commonly associated proteins with EFR, FLS2, and RBOHD in *Arabidopsis thaliana*. A) Comparison of candidate-associated proteins with EFR, FLS2, and RBOHD identified by co-immunoprecipitation. The Venn diagram illustrates candidate-associated proteins identified by IP of EFR-GFP, FLS2-GFP or 3xFLAG-RBOHD from Arabidopsis seedlings of *efr-1/pEFR:EFR-GFP* (Kadota et al., 2014), *fpFLS2:FLS2-GFP*, or *rbohD/pRBOHD:3×FLAG-gRBOHD* (Goto et al., 2023). The protein list is shown in Supplemental Data Set S1. **B)** The list of commonly associated proteins with EFR, FLS2, and RBOHD. An asterisk indicates the known components of the PRR complex, and the double asterisks indicate proteins accumulate in detergent-resistant membrane compartments in response to flg22 (Keinath et al., 2010).

QSK1 (AT3G02880) is of particular significance as multiple tryptic peptides could be identified in the IPs with FLS2, EFR, and RBOHD (Supplemental Data Set S2). Notably, transient expression of *QSK1-3xHA* in *Nicotiana benthamiana* led to significant reduction in flg22-induced ROS production (Goto et al., 2023) (Supplemental Fig. S1). QSK1 is an LRR-RK with five LRRs in its ectodomain (Isner et al., 2018; Wu et al., 2019). To independently validate the interaction of QSK1 with FLS2, EFR, and RBOHD in Arabidopsis, we generated α-QSK1 antibodies. IP of FLS2-GFP from the *pFLS2:FLS2-GFP* stable transgenic line revealed a clear ligand-independent association between FLS2-GFP and endogenous QSK1 (Fig. 2A), in contrast to the ligand-dependent FLS2-BAK1 interaction. Further, we conducted IP experiments with EFR-GFP and 3xFLAG-RBOHD from *efr-1/pEFR:EFR-GFP* and *rbohD/pRBOHD:3xFLAG-RBOHD*, respectively (Fig. 2, B to C). The data reveal that RBOHD and EFR form ligand-independent interactions with QSK1, suggesting that QSK1 is an integral component of the PRR-RBOHD complex prior to elicitation, and this association remains stable even after PAMP treatment.

**Figure 2.**
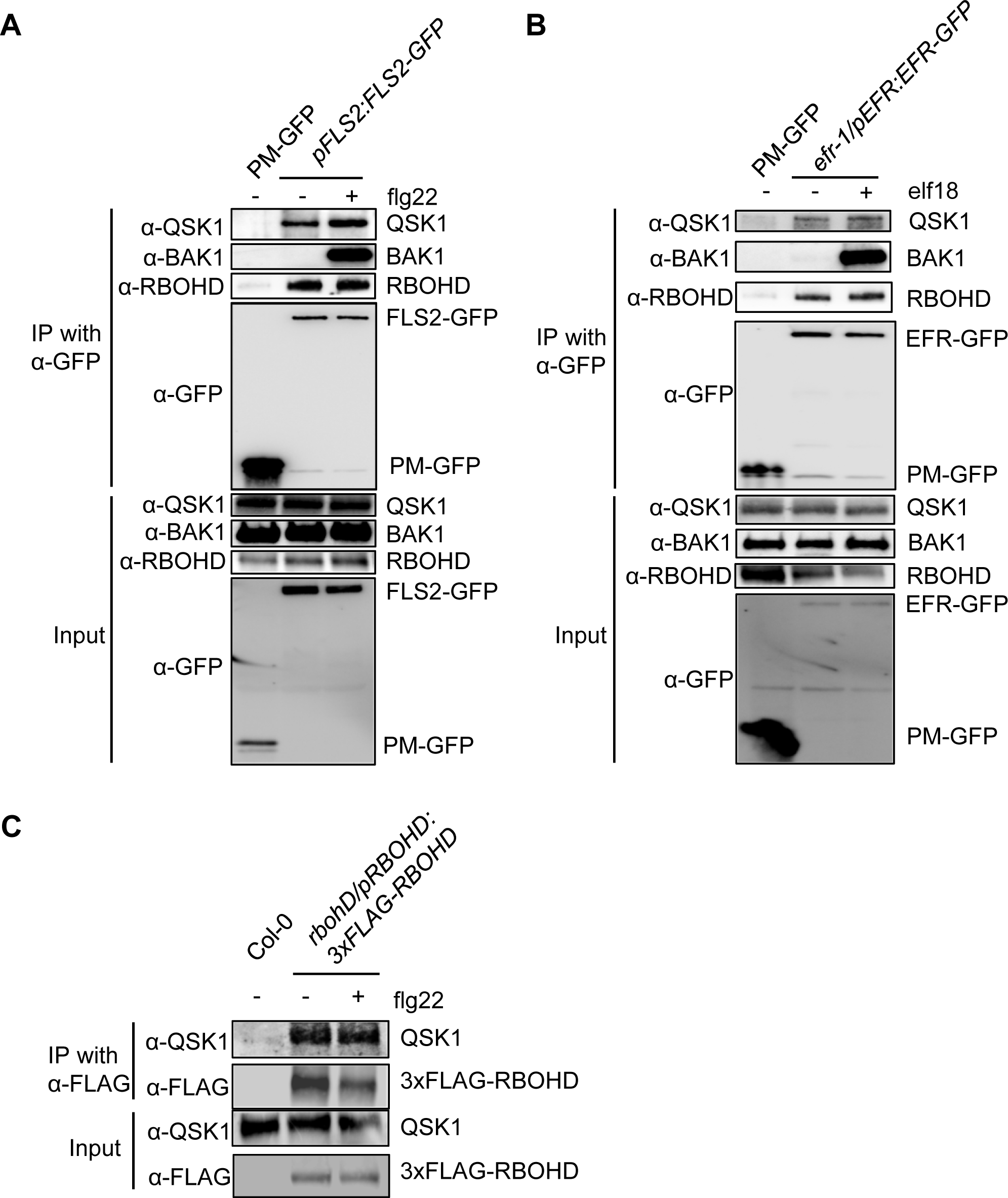
QSK1 forms a stable complex with FLS2, EFR, and RBOHD in Arabidopsis thaliana. **A and B)** Two-week-old Arabidopsis seedlings of *pFLS2:FLS2-GFP*, *efr-1/pEFR:EFR-GFP*, or PM-GFP (*p35S: LTI6b-GFP*) were treated with or without 1 µM flg22 or 1 µM flg22 for 10 min. Total proteins (input) were immunoprecipitated with α-GFP magnetic beads, followed by immunoblotting with α-GFP, α- QSK1, α-BAK1, and α-RBOHD antibodies. LTI6b, a known plasma membrane protein was used as a control to illustrate that QSK1, RBOHD, and BAK1 do not associate with GFP at the plasma membrane. **C)** Two-week-old Arabidopsis seedlings of rbohD/pRBOHD:3xFLAG-RBOHD or Col-0 were treated with or without 1 µM flg22 for 10 min, and the total proteins were immunoprecipitated with α-FLAG magnetic beads followed by immunoblotting with α-FLAG and α-QSK1 antibodies. Col-0 plants were used as a control to illustrate that QSK1 does not associate with α-FLAG nonspecifically. All the experiments were repeated three times with similar results.

### QSK1 negatively regulates PTI

To elucidate the role of QSK1 in the regulation of PRR-RBOHD complexes, we conducted comprehensive characterization of the Arabidopsis *qsk1* mutant (SALK_ 019840) (Isner et al., 2018). The *qsk1* mutant harbors a T-DNA insertion within the first exon, resulting in pronounced reduction in *QSK1* transcript levels compared to Col-0 (Supplemental Fig. S2, A to B). In addition, immunoblotting with α-QSK1 antibodies failed to detect the QSK1 protein in the *qsk1* mutant (Supplemental Fig. S2C), indicating that *qsk1* is a null mutant. The *qsk1* mutant exhibited a significant increase in ROS production in response to flg22, elf18, and the DAMP peptide pep1 (Figs 3, A to B; Supplemental Fig. S2D). Furthermore, this mutant also showed enhanced MAPK activation 15 minutes following flg22 treatment (Fig. 3C). Collectively, these results indicate that QSK1 exerts a negative regulatory influence on PTI signaling pathways.

**Figure 3.**
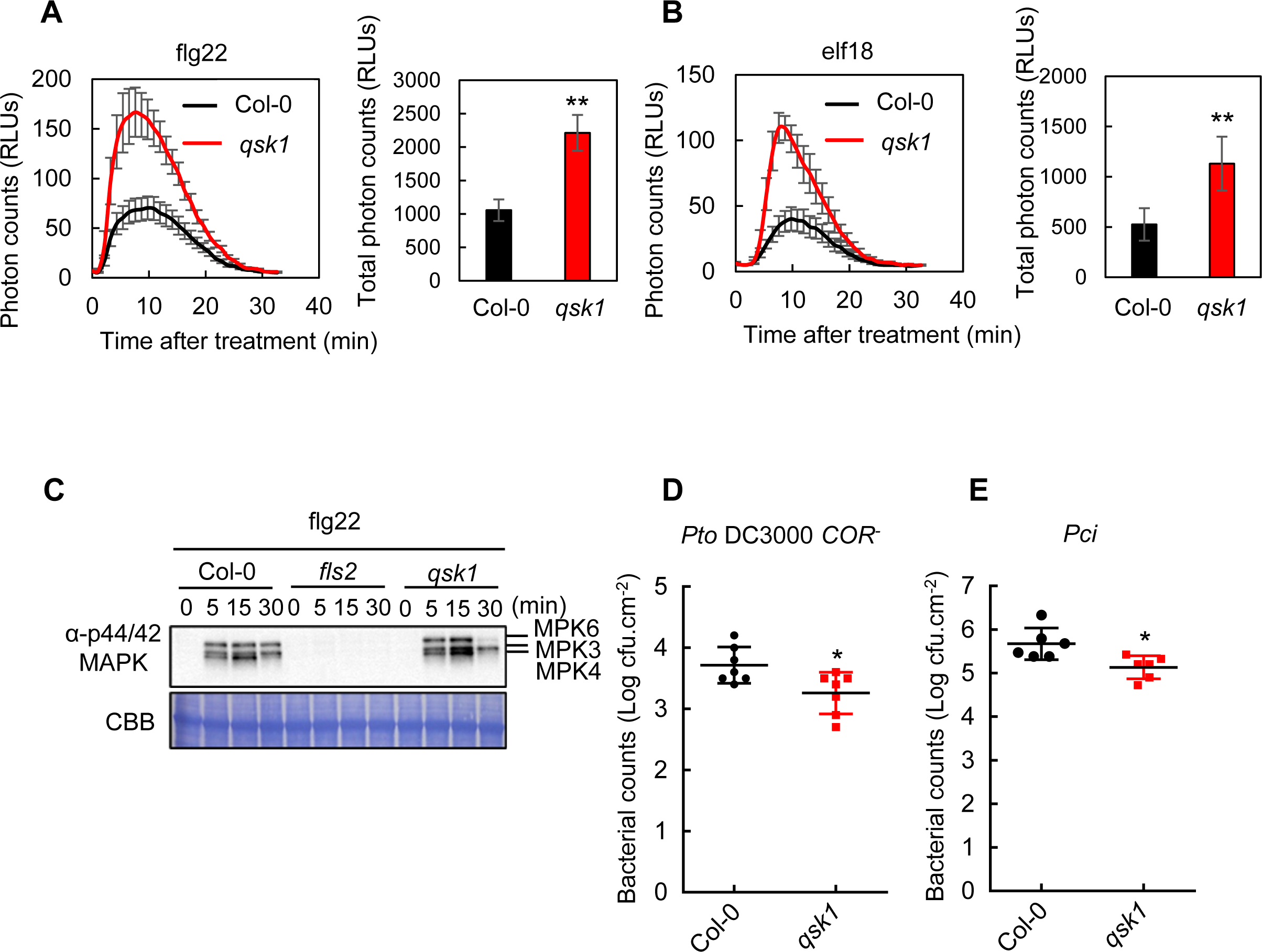
Arabidopsis *qsk1* mutant shows enhanced PTI responses compared to Col-0. **A and B)** *qsk1* mutant has enhanced ROS production following treatment with flg22 and elf18. Eight leaf discs from four- to five-week-old Arabidopsis plants were treated with 1 µM flg22 (**A**) or 1 µM elf18 (**B**), and time-course (left) and the total amount (right) of ROS production was measured by a luminol-based assay. Values are mean ± standard error (SE) (n=8). An asterisk indicates significant differences (Student’s t-test, \**p* ≤ 0.05). **C)** *qsk1* mutant induced enhanced MAPKs activation following treatment with flg22. Ten-day-old Arabidopsis seedlings were treated with 1 µM flg22 and phosphorylated MAPKs were detected on immunoblotting with α-phospho-p44/42 MAPK (Erk1/2) (Thr202/Tyr204) antibody. Equal loading of protein samples is shown by Coomassie Brilliant Blue (CBB) staining. **D and E**) *qsk1* mutant was more resistant to bacteria. *Pseudomonas syringae* pv. *tomato* (*Pto*) DC3000 lacking the toxin coronatine (*COR^-^*) (**D**) or *Pseudomonas syringae* pv. *cilantro* (*Pci*) 0788-9 (**E**) were sprayed onto leaf surfaces of six-week-old soil-grown Arabidopsis plants at a concentration of 1x10^5^ cfu (colony-forming units)/mL. Three-day post spray-inoculation, leaves were harvested to determine bacterial growth. Data are means ± SE of 6 replicates. An asterisk indicates significant differences (Student’s t-test, \**p* ≤ 0.05). All the experiments were repeated three times with similar results.

To gain further insights into the impact of QSK1 on disease resistance, we assessed growth of the weakly virulent bacterial strain *Pto* DC3000 *COR^-^* which lacks the toxin coronatine (COR) responsible for inducing stomatal reopening during infection (Melotto et al., 2006), and the non-adapted bacterium *Pseudomonas syringae* pv. *Cilantro* (*Pci*) 0788-9, known to exhibit poor growth on Col-0 plants (Lewis et al., 2008). Six-week-old Arabidopsis plants were spray-inoculated with *Pto* DC3000 *COR^-^*and *Pci.* At three days post-inoculation (dpi), *qsk1* demonstrated enhanced resistance compared to Col-0 (Fig. 3, D to E). This highlights the significant role of QSK1 in the negative regulation of plant resistance to bacterial disease.

To verify that the observed phenotype is due to the lack of *QSK1*, we generated the complementation line, *qsk1/pQSK1:QSK1-GFP*. This complementation reversed the enhanced PAMP-induced ROS production evident in the *qsk1* mutant (Supplemental Fig. S3, A to B). No morphological differences were observed among the *qsk1* mutant, *qsk1/pQSK1:QSK1-GFP* lines, and Col-0 (Supplemental Fig. S3C). These results confirm that the amplified PTI responses in the *qsk1* mutant are attributed to the absence of *QSK1*.

To further investigate the role of QSK1 in modulating PRR-RBOHD complexes, we generated two independent Arabidopsis transgenic lines overexpressing *QSK1-3×HA* under the control of the *CaMV 35S* promoter (*p35S:QSK1-3×HA*). These lines exhibited markedly elevated *QSK1* transcript levels compared to Col-0 (Supplemental Fig. S4A) and produced a significantly higher amount of QSK1-3xHA protein than the endogenous QSK1 (Supplemental Fig. S4B). Morphological evaluations highlighted that the *p35S:QSK1-3×HA* lines had a marginally reduced size compared to both Col-0 and the *qsk1* mutant (Supplemental Fig. S4C). In stark contrast to the *qsk1* mutant, the *p35S:QSK1-3×HA* lines exhibited notably diminished ROS production upon treatment with flg22 and elf18 in comparison to Col-0 (Fig. 4, A to B). Additionally, *p35S:QSK1-3×HA* lines displayed attenuated MAPK activation in response to flg22 (Fig. 4C) and showed reduced resistance to *Pto* DC3000 *COR^-^* and *Pci* compared to Col-0 (Fig. 4, D to E). These results confirm that *QSK1* plays an important role as a negative regulator in PTI in Arabidopsis.

**Figure 4.**
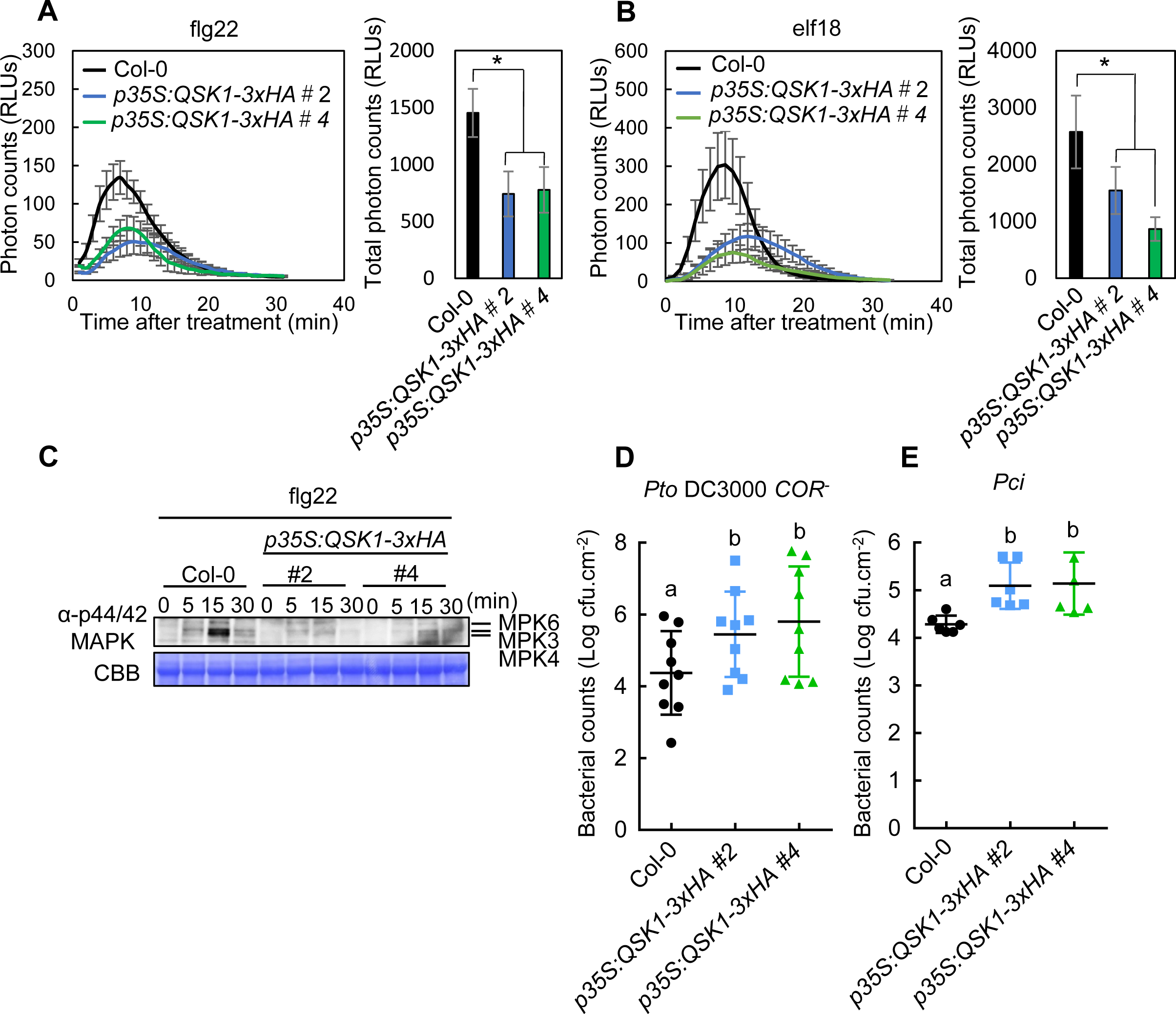
Arabidopsis *QSK1* over expression lines (*p35S:QSK1×HA* QSK1) have reduced PTI responses compared to Col-0. **A and B)** *p35S:QSK1×HA* lines showed reduced ROS production in response to flg22 and elf18. Eight seven-day-old Arabidopsis seedlings were treated with 1 µM flg22 (**A**) or 1 µM elf18 (**B**), and time-course (left) and the total amount (right) of ROS production was measured by a luminol-based assay. Values are mean ± SE (n=8). An asterisk indicates significant differences (Student’s t-test, \**p* ≤ 0.05). **C)** *p35S:QSK1×HA* lines showed reduced MAPKs activation in response to flg22. Ten-day-old Arabidopsis seedlings were treated with 1 µM flg22 and phosphorylated MAPKs were detected on immunoblotting with α-phospho-p44/42 MAPK (Erk1/2) (Thr202/Tyr204) antibody. Equal loading of protein samples is shown by CBB staining. **D and E)** *p35S:QSK1×HA* lines were more susceptible to bacteria. *Pto* DC3000 C*OR*^-^ (**D**) or *Pci* (**E**) were sprayed onto leaf surfaces of six-week-old soil-grown Arabidopsis plants at a concentration of 1x10^5^ cfu/mL. Three-day post spray-inoculation, leaves were harvested to determine bacterial growth. Data are means ± SE of 6 replicates. Different letters indicate significantly different values at *p* ≤ 0.05 (one-way ANOVA, Tukey post hoc test). All the experiments were repeated three times with similar results.

To determine the subcellular localization of QSK1 in plant cells, we transiently expressed a QSK1-GFP fusion protein in *N. benthamiana*. QSK1-GFP localizes at the plasma membrane (Supplemental Fig. S5, A to B). This subcellular localization was confirmed in Arabidopsis using a stable transgenic line, *qsk1*/*pQSK1:QSK1-GFP* (Supplemental Fig. S5C). Additionally, we examined the transcriptional response of *QSK1* to PAMPs. Treatment with flg22 and elf18 led to an increase in *QSK1* transcript levels, indicating its transcriptional upregulation upon PAMP recognition (Supplemental Fig. S6).

### QSK1 negatively regulates PRR protein levels

Since QSK1 negatively regulates both ROS production and MAPK activation, two distinct signaling events following PAMP recognition (Xu et al., 2014), we hypothesized that QSK1 might influence the activity or stability of PRRs. Immunoblotting showed elevated FLS2 protein abundance in the *qsk1* mutant relative to Col-0 and the complemented *qsk1*/*pQSK1:QSK1-GFP* lines, while BAK1 levels remained unaffected (Fig. 5A). Conversely, FLS2 protein levels were reduced in QSK1 overexpression lines (*p35S:QSK1-3×HA*) compared to Col-0 (Fig. 5B). This regulatory mechanism does not appear to operate at the transcriptional level since *FLS2* mRNA amounts were comparable among Col-0, *qsk1*, and *p35S:QSK1-3×HA lines* (Fig. 5C). Supporting this notion, *N. benthamiana* plants co-expressing *FLS2-GFP* and *QSK1-GFP* under the control of the p35S promoters exhibited reduced FLS2-GFP protein levels (Fig. 5D). Similarly, overexpression of *QSK1* led to a decline in EFR protein levels; the EFR-GFP levels in *pEFR:EFR-GFP/ p35S:QSK1-3×HA* line were lower than those in *pEFR:EFR-GFP* lines (Fig. 5E). Further investigation into the impact of QSK1 on the subcellular distribution of FLS2-GFP, revealed that a notable reduction in plasma membrane localization when co-expressed with *QSK1* in the *pFLS2:FLS2-GFP*/*p35S:QSK1-3xHA* line (Fig. 5F). These results suggest that QSK1 exerts a negative regulatory effect on PRR protein accumulation at the plasma membrane.

**Figure 5.**
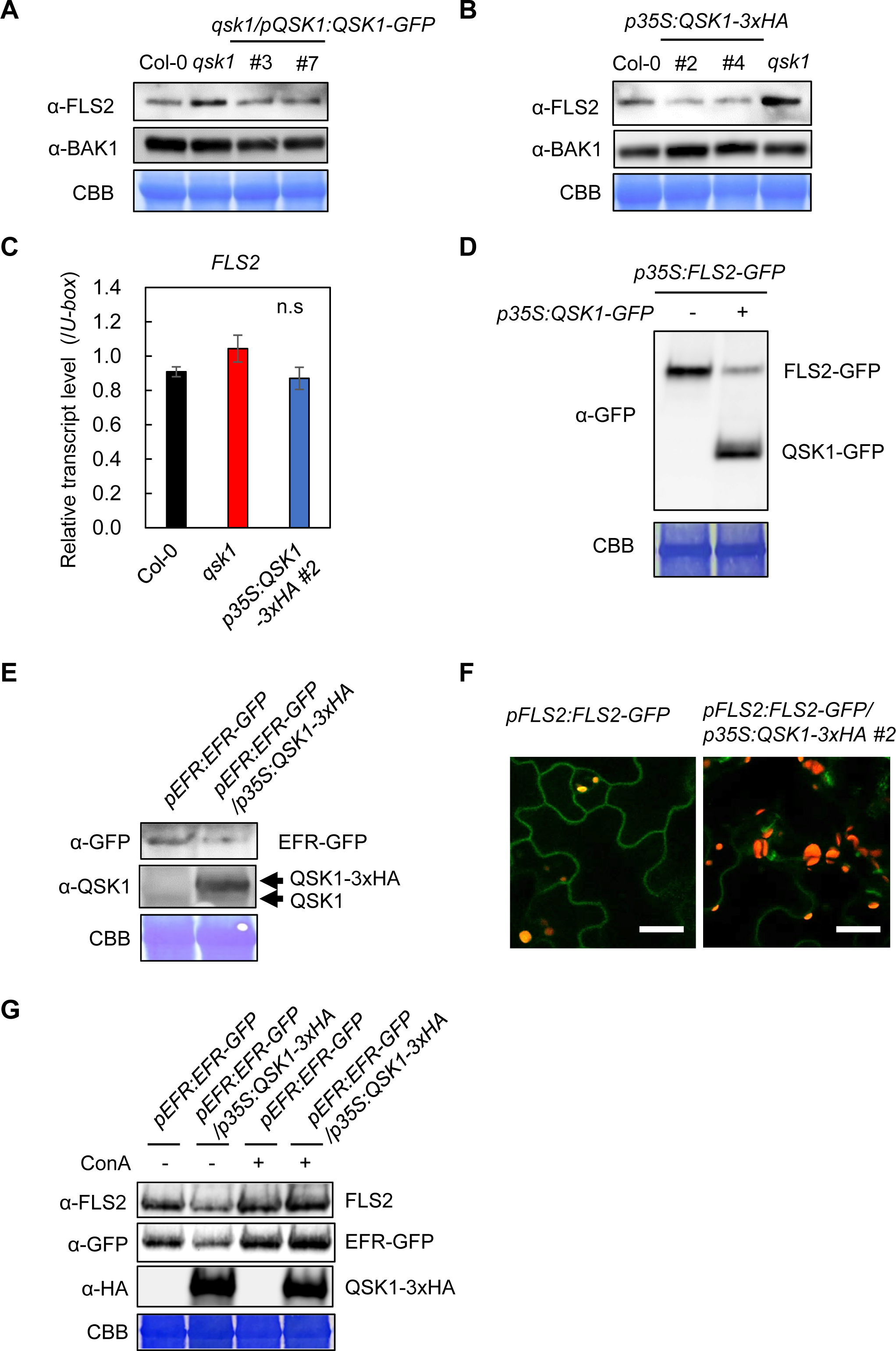
QSK1 negatively regulates FLS2 and EFR accumulation. **A)** FLS2 protein accumulates more in *qsk1* mutant than in Col-0 and the complementation lines (*qsk1/pQSK1:QSK1-GFP*). **B)** FLS2 protein accumulates less in *p35S:QSK1×HA* lines than in Col-0. FLS2 and BAK1 protein levels of two-week-old Arabidopsis seedlings were measured by immunoblotting with α-FLS2 and α- BAK1 antibodies. Equal loading of protein samples is shown by CBB staining. **C)** FLS2 transcript levels are not changed in Col-0, *qsk1* mutant, and *p35S:QSK1×HA* lines. Transcript levels of *FLS2* in two-week-old Arabidopsis seedlings were measured by RT-qPCR after normalization to the *U*-*box* housekeeping gene transcript (*At5g15400*). Values are mean ± SE of three biological replicates. There are no significant differences at *p* ≤ 0.05 (one-way ANOVA, Turkey’s *post hoc* test). **D**) The expression of *QSK1-GFP* reduces FLS2-GFP protein levels in *Nicotiana benthamiana.* FLS2-GFP and QSK1-GFP proteins were transiently expressed under the control of *p35S* promoter and their protein levels were measured three days after agroinfiltration by immunoblotting with α-GFP antibodies. Agrobacterium concentration (OD600=0.6) was adjusted with empty Agrobacterium. **E)** QSK1 reduces EFR protein levels. Protein levels of EFR-GFP and QSK1 in two-week-old Arabidopsis seedlings of *pEFR:EFR-GFP* and *pEFR:EFR-GFP*/*p35S:QSK1-3xHA* were measured by immunoblotting with α-GFP antibodies. **F)** QSK1 reduces FLS2 protein accumulation at the plasma membrane. The localization of FLS2-GFP in cotyledons of ten-day-old seedlings of *pFLS2:FLS2-GFP* line and *pFLS2:FLS2-GFP/p35S:QSK1-3xHA#2* line were observed by confocal microscopy. The white bars represent 30 μm. **G)** Concanamycin A (ConA) suppresses QSK1-mediated PRR reduction. Two-week-old Arabidopsis seedlings of *pEFR:EFR-GFP* and *p35S:QSK1-3xHA*/*pEFR:EFR-GFP* lines were treated with or without 1 μM ConA for 10 h. The protein levels of FLS2, EFR-GFP, and QSK1-3xHA were measured by immunoblotting. Equal loading of protein samples is shown by CBB staining. All the experiments were repeated three times with similar results.

To elucidate the mechanism behind QSK1’s modulation of FLS2 protein levels, we employed a pharmacological approach, using an array of inhibitors: MG132 (proteasome x), Bafilomycin A1 (vacuolar-type-H^+^-ATPase inhibitor), E-64d (cysteine protease inhibitor), TLCK (serine protease inhibitor), Wortmannin (phosphatidylinositol 3-kinase inhibitor), Brefeldin A (ER-Golgi transport inhibitor), Cycloheximide (protein synthesis inhibitor), and Concanamycin A (ConA, vacuolar-type-H^+^-ATPase inhibitor) (Fig. 5G; Supplemental Fig. S7). Notably, ConA mitigated the QSK1-mediated reduction of both FLS2 and EFR levels (Fig. 5G). ConA is known to block vacuolar transport, thereby impeding autophagic degradation pathway as well as the endocytosis-mediated degradation pathway (Dettmer et al., 2006; Scheuring et al., 2011). These findings suggest that *QSK1* overexpression may facilitate vacuolar degradation of PRRs through the autophagy pathway or the endocytosis pathway.

### HopF2*_Pto_*-HA interacts with QSK1 and reduces FLS2 protein levels

QSK1 could represent a potential effector target as part of PRR complexes because plant pathogens often deploy virulence effectors to target the PRR complex to effectively suppress PTI. Our attention was drawn to HopF2*_Pto_* from *Pto* DC3000, renowned for its potent inhibition of early PTI responses (Wilton et al., 2010; Wu et al., 2011; Hurley et al., 2014; Zhou et al., 2014), as a likely candidate effector targeting QSK1, for several reasons. Firstly, Khan et al., conducted enzyme-catalyzed proximity labeling of HopF2*_Pto_* (Proximity-dependent Biotin Identification (BioID)) (Khan et al., 2018) and identified QSK1 as one of the 19 biotinylated proteins. Secondly, we employed a combination of yeast two-hybrid methods with next-generation sequencing, known as QIS-seq (Lewis et al., 2012), and revealed QSK1 as one of the 15 potential targets (Fig. 6A; Supplemental Data Set S3_1). Thirdly, a comparative analysis of potential HopF2*_Pto_* interactors by QIS-seq (Quantitative Interactor Screening with Next-Generation Sequencing) and BioID, alongside PRR complex components, using a Venn diagram (Fig. 6A), highlighted QSK1 as the sole common factor across all three datasets (Fig. 6A; Supplemental Data Set S3_2). This finding aligns with previous IP-MS experiments by Hurley *et al*., which also listed QSK1 among the proteins interacting with HopF2*_Pto_* when expressed in Arabidopsis (Hurley et al., 2014).

**Figure 6.**
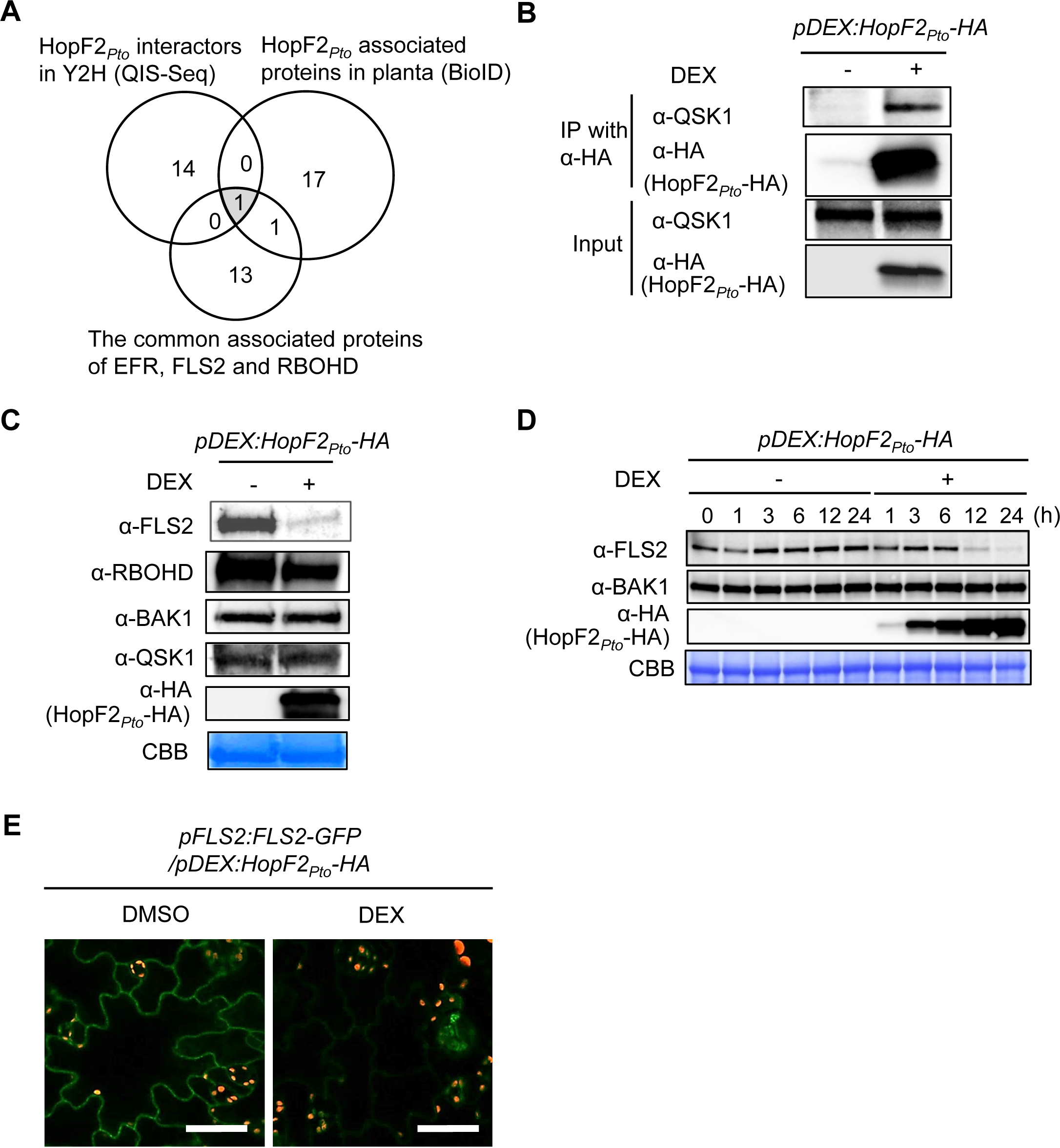
HopF2*_Pto_* associates with QSK1 and reduces FLS2 protein level. **A)** Comparison of candidate interactors of HopF2*_Pto_* and the commonly associated proteins with EFR, FLS2, and RBOHD. The Venn diagram illustrates candidate HopF2*_Pto_* interactors identified by yeast two-hybrid screening coupled with next-generation sequencing (QIS-Seq) and by proximity-dependent biotin identification (BioID) in planta (Khan et al., 2018) with the commonly associated proteins with EFR, FLS2, and RBOHD identified in this study. **B)** HopF2*_Pto_* associates with QSK1 *in vivo*. Two-week-old Arabidopsis seedlings of *pDEX:HopF2_Pto_-HA* were treated with or without 30 µM dexamethasone (DEX) for 24 h. Total proteins (input) were immunoprecipitated with α-HA magnetic beads followed by immunoblotting with α-HA and α-QSK antibodies. **C and D**) HopF2*_Pto_* specifically reduced FLS2 protein accumulation. Two-week-old Arabidopsis seedlings of *pDEX:HopF2_Pto_-HA* were treated with or without 30 µM DEX and FLS2, RBOHD, BAK1, QSK1, and HopF2*_Pto_*-HA protein levels were measured by immunoblotting. Equal loading of protein samples is shown by CBB staining. **E)** HopF2*_Pto_* reduced FLS2 protein accumulation at the plasma membrane. Ten-day-old seedlings of *pFLS2:FLS2-GFP/ pDEX:HopF2_Pto_-HA* line were treated with or without 30 µM DEX for 24 h and the localization of FLS2-GFP in cotyledons was observed by confocal microscopy. The white bar represents 50 μm. All the experiments were repeated three times with similar results.

To validate the interaction between HopF2*_Pto_*-HA and endogenous QSK1 *in vivo*, we employed the dexamethasone (DEX)-inducible system in transgenic Arabidopsis carrying the *pDEX:HopF2_Pto_-HA* construct. Our results show *in vivo* interaction between HopF2*_Pto_*-HA and QSK1 upon DEX treatment (Fig. 6B). To assess the impact of HopF2*_Pto_*on PRR complexes, we examined the protein levels of FLS2, RBOHD, BAK1, and QSK1 with or without expression of *HopF2_Pto_-HA* (Fig. 6C). Strikingly, HopF2*_Pto_*-HA specifically diminished the protein levels of FLS2 without affecting the other proteins. The reduction in FLS2 coincided with an increase in the levels of HopF2*_Pto_*-HA following DEX treatment (Fig. 6D). Next, we examined the effects of HopF2*_Pto_*-HA on the subcellular localization of FLS2-GFP (Fig. 6E). In the absence of HopF2*_Pto_*-HA expression, FLS2-GFP predominantly localized to the plasma membrane. However, induction of *HopF2_Pto_-HA* expression by DEX treatment led to a significant reduction of FLS2-GFP at the plasma membrane.

### The catalytic residue D175 of HopF2*_Pto_* is required for its virulence function

A mutation in the catalytic residue D175 (D175A) of HopF2*_Pto_* leads to a significant reduction of its virulence, indicating the indispensable role of mono-ADP ribosylation (MARylation) in the functionality of HopF2*_Pto_* (Wilton et al., 2010). Notably, DEX-induced expression of *HopF2_Pto_(D175A)-HA* did not decrease FLS2 protein levels (Fig. 7A), suggesting that MARylation activity is essential for HopF2*_Pto_*’s ability to deplete FLS2. To further investigate the effects of HopF2*_Pto_* and its MARylation activity on FLS2 during infection, we introduced both the wild-type HopF2*_Pto_*-HA and its D175A mutant into the non-pathogenic bacteria *Pseudomonas fluorescens* Pf0-1 (Fig. 7B). We selected *P. fluorescens* Pf0-1 due to its absence of virulence effectors, allowing a focused examination of HopF2*_Pto_* effects. The *bak1-5 bkk1* double mutants were infected with these modified bacteria. Employing *bak1-5 bkk1* mutants aimed to minimize the PTI-induced FLS2 accumulation during infection, although the suppression of FLS2 accumulation after bacterial inoculation was not complete. Infection with *P. fluorescens* Pf0-1 harboring *HopF2_Pto_-HA* for ten hours resulted in increased levels of HopF2*_Pto_*-HA and a concurrent decrease in FLS2 levels, compared to both untransformed *P. fluorescens* Pf0-1 and *P. fluorescen*s Pf0-1 harboring *HopF2_Pto_*(D175A)*-HA*. These data demonstrate that HopF2*_Pto_*-HA actively reduces FLS2 protein levels during infection and that the MARylation activity of HopF2*_Pto_*is required for this function. A pharmacological assay that involved a range of inhibitors, including ConA, E-64d, 3-Methyladenine (3-MA, PI3K inhibitor), BAF, Wm, BFA, MG132, and TLCK, showed that none of these inhibitors succeeded in counteracting the FLS2 depletion induced by HopF2*_Pto_*(Supplemental Fig. S8).

**Figure 7.**
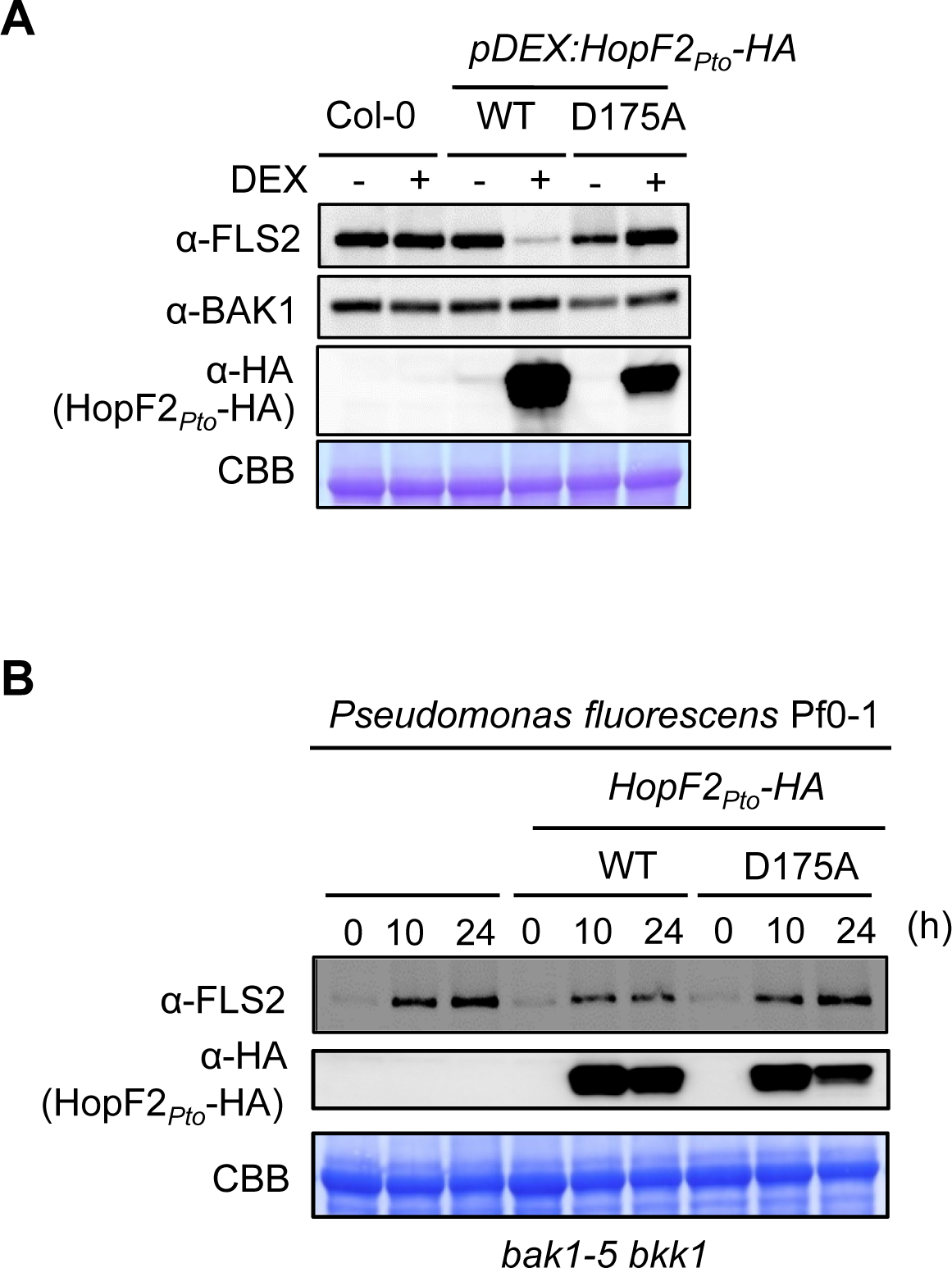
Mono ADP ribosylation (MARylation) activity of HopF2*_Pto_* is required for the FLS2 elimination. **A)** The catalytic residue D175 for MARylation activity in HopF2_Pto_ is required for the inhibition of FLS2 accumulation. Two-week-old Arabidopsis seedlings of *pDEX: HopF2_Pto_-HA* and *pDEX:HopF2_Pto_* (D175A)*-HA* were treated with 30 µM DEX for 24 h and FLS2, BAK1, and HopF2*_Pto_* -HA protein levels were measured by immunoblotting. **B)** HopF2*_Pto_* inhibits FLS2 protein accumulation during infection. Immunoblotting detecting FLS2 and HopF2*_Pto_*-HA in Col-0 during bacterial infection after syringe inoculation with *Pseudomonas fluorescens* Pf0-1, *P. fluorescens* Pf0-1 *HopF2_Pto_-HA*, or *P. fluorescens* Pf0-1 *HopF2_Pto_* (D175A)*-HA*. All the experiments were repeated three times with similar results.

### HopF2*_Pto_* modulates the expression of immune-related genes in Arabidopsis

To explore the influence of HopF2*_Pto_*on plant immune responses, RNA-seq analysis was performed on Arabidopsis Col-0 and *pDEX:HopF2_Pto_-HA* seedlings, post 24-h treatment with either DMSO or DEX. The multidimensional scaling plot displayed consistent global gene expression patterns across all four biological replicates for both treatments (Supplemental Fig. S9A). Notably, the *pDEX:HopF2_Pto_-HA* line exhibited significant transcriptional changes upon DEX treatment, whereas DEX treatment in Col-0 led to only minor alterations in gene expression compared to those in the Col-0 and *pDEX:HopF2_Pto_-HA* lines treated with DMSO.

To differentiate gene expression changes induced by HopF2*_Pto_* from those solely caused by DEX, we compared the gene expression in the DEX-treated *pDEX:HopF2_Pto_-HA* line with DEX-treated Col-0. In the DEX-treated *pDEX:HopF2_Pto_-HA* line, we observed an upregulation of 1399 genes and a downregulation of 2,708 genes by at least twofold, along with 330 genes upregulated and 879 genes downregulated by at least fourfold (Supplemental Data Set S4). Gene Ontology (GO) enrichment analyses conducted on highly upregulated (330 genes, log2 fold change ≥ 2, FDR ≤ 0.05) and highly downregulated (879 genes, log2 fold change ≤ −2, FDR ≤ 0.05) genes provided insights into the biological significance of these transcriptional changes (Supplemental Data Set S5_1 and S5_2). Remarkably, both upregulated and downregulated genes were significantly associated with GO terms related to biotic stress responses and immunity, underlining HopF2*_Pto_*’s crucial role in modulating specific immune-related genes in Arabidopsis.

To pinpoint genes distinctively affected by *HopF2_Pto_*expression, self-organizing map (SOM) clustering was applied to the most differentially expressed genes, focusing on the top 25% based on their coefficient of variation across samples. These genes were grouped into 12 clusters, reflecting unique expression patterns in Col-0 and *pDEX:HopF2_Pto_-HA* following either DMSO or DEX treatment (Supplemental Fig. S9B; Supplemental Data Set S6). Notably, genes in cluster 1 were exclusively upregulated by HopF2*_Pto_*, whereas those in cluster 2 were specifically downregulated. The GO enrichment analysis revealed that both clusters were enriched in GO terms associated with biotic stress responses and immunity (Supplemental Data Set S5_3 and S5_4), and cluster 2 exhibited a pronounced enrichment for GO terms like “membrane”, “cell periphery”, and “plasma membrane”. These observations suggest that HopF2*_Pto_* selectively modulates gene expression related to immune response and plasma membrane-associated proteins.

Given HopF2*_Pto_*’s role in diminishing FLS2 levels, we assessed the transcript levels of known *PRRs* (Fig. 8A). Notably, our data showed that HopF2*_Pto_* significantly reduces the transcript levels of certain *PRRs*, such as *FLS2*, *LORE* (*LIPOOLIGOSACCHARIDE-SPECIFIC REDUCED ELICITATION,* a PRR for bacterial fatty acid metabolite 3-OH-C10:0)(Kutschera et al., 2019), and *MIK2* (*MALE DISCOVERER 1-INTERACTING RECEPTOR-LIKE KINASE 2*, a PRR for SCOOP phytocytokines) (Hou et al., 2021; Rhodes et al., 2021), as well as *IOS1*, an import regulator in PRR complexes (Yeh et al., 2016) (Supplemental Fig. S10). Such reduction in transcript levels likely contributes to HopF2*_Pto_*’s suppression of PTI responses, as corroborated by our observation that HopF2*_Pto_*inhibits ROS production mediated by FLS2 and MIK2 (Supplemental Fig. S11).

**Figure 8.**
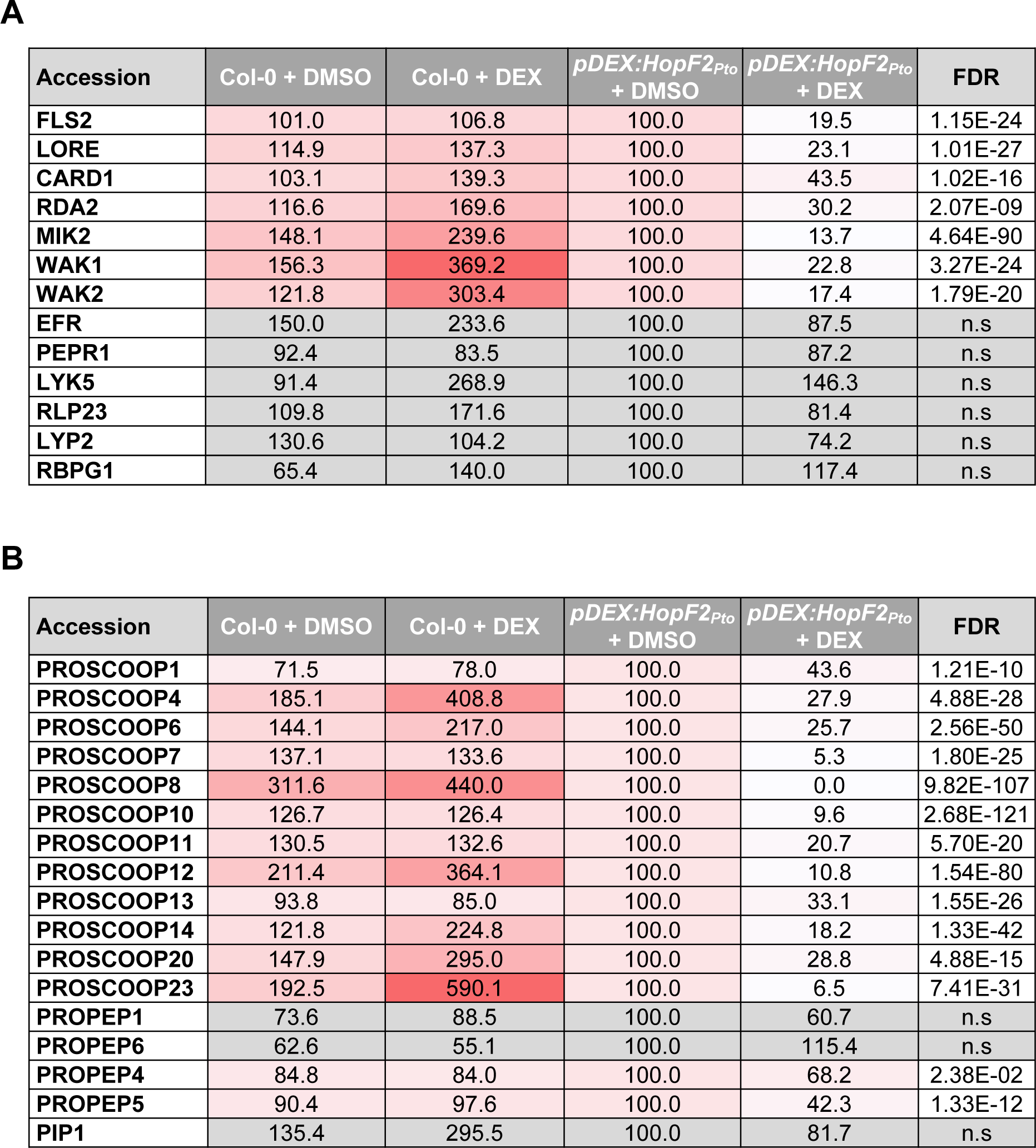
HopF2*_Pto_* reduces transcript levels of *PRRs* and *PROSCOOPs*. Transcript levels of *PRRs* (**A**), *PROSCOOPs*, *PROPEPs,* and *PIP1* (**B**) were measured by RNA-seq in two-week-old seedlings of Col-0 and *pDEX:HopF2_Pto_-HA* after treatment with 30 µM DEX for 24 h. The relative expression values of the genes are shown compared to the “*pDEX:HopF2_Pto_* + DMSO” control. The FDR values between “*pDEX:HopF2_Pto_* + DMSO” and “*pDEX:HopF2_Pto_* + DEX” are shown. Grey boxes in the heat map indicate no statistically significant difference at FDR ≤ 0.05.

### HopF2*_Pto_* reduces transcript levels of most PROSCOOPs

Beyond inhibiting *PRR* gene expression, HopF2*_Pto_* also downregulates the SCOOP phytocytokine signaling. SCOOP phytocytokines, exclusive to the Brassicaceae family, are a unique group of peptides that are cleaved from the C-terminus of their respective precursors, termed PROSCOOPs (Gully et al., 2019). Our transcriptomic analysis revealed that HopF2*_Pto_* significantly downregulates the transcript levels of multiple *PROSCOOPs*, especially *PROSCOOP7*, *8*, *10*, *12*, and *23* (Fig. 8B), while its effect on *PROPEPs* and *PROPIP1*, encoding other DAMP peptides is minimal. This suggests HopF2*_Pto_*’s role in attenuating SCOOP phytocytokine signaling by downregulating both *PROSCOOPs* and *MIK2* gene expression.

### HopF2*_Pto_* reduces EFR protein levels possibly through vacuolar degradation

While HopF2*_Pto_* reduces the expression of *FLS2*, *LORE*, and *MIK2*, it does not affect the expression of other *PRRs* such as *EFR* and *PEPR1* (*PEP RECEPTOR1,* a PRR for Pep1 and Pep2 peptides) (Fig. 8A). Nevertheless, HopF2*_Pto_* effectively impairs ROS production and MAPK activation triggered by these PRRs (Supplemental Figs. S11 and S12), indicating that HopF2*_Pto_* may also employ a transcription-independent mechanism to inhibit PTI. This insight prompted further exploration into how HopF2*_Pto_* affects the EFR signaling pathway. We generated a homozygous *pDEX:HopF2_Pto_-HA/pEFR:EFR-GFP* line to assess the impact of *HopF2_Pto_-HA* expression on EFR-GFP levels. Remarkably, DEX-induced *HopF2_Pto_-HA* expression led to a decrease in EFR protein levels, suggesting that HopF2*_Pto_*exerts its influence on EFR protein levels via transcription-independent mechanisms. Interestingly, ConA effectively countered the HopF2*_Pto_*-mediated reduction in EFR protein levels (Fig. 9), implying that this reduction might occur via vacuolar degradation through either the autophagy pathway or the endocytosis pathway. Additionally, we assessed the effect of the proteasome inhibitor MG132, which only slightly inhibited the reduction in EFR protein levels.

**Figure 9.**
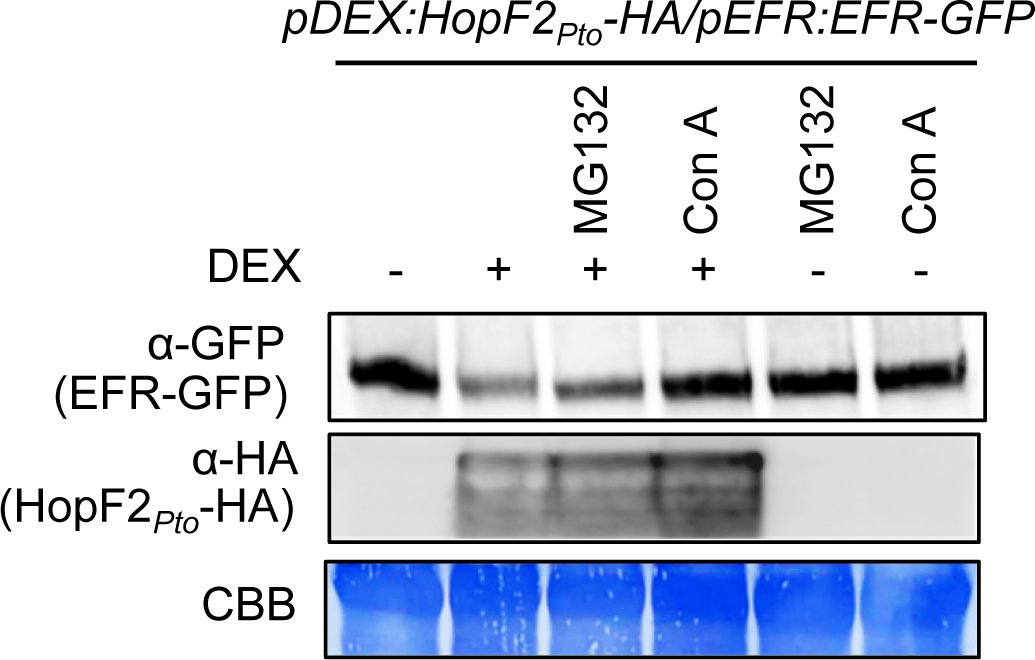
Con A inhibits HopF2*_Pto_* -mediated reduction of EFR expression. Two-week-old Arabidopsis seedlings of *pDEX:HopF2_Pto_-HA*/*pEFR:EFR-GFP* were treated with or without 30 µM DEX for 24 h, followed by the treatment with DMSO, 100 μM MG132, or 1 μM ConA for 10 h. The protein levels of EFR-GFP and HopF2*_Pto_*-HA were measured by immunoblotting with α-GFP and α-HA antibodies. Equal loading of protein samples is shown by CBB staining. This experiment was repeated three times with similar results.

### HopF2*_Pto_* requires QSK1 for its stabilization

To investigate the functional relationship between HopF2*_Pto_* and QSK1, we generated a *qsk1/ pDEX:HopF2_Pto_-HA* homozygous line by crossing, and checked the HopF2*_Pto_*-mediated reduction of FLS2 protein in a *qsk1* knockout background (Fig. 10A). Remarkably, the absence of QSK1 significantly reduces HopF2*_Pto_*’s ability to reduce FLS2 levels, showing the crucial role of QSK1 in HopF2*_Pto_* function. Intriguingly, HopF2*_Pto_*-HA protein levels were decreased in the *qsk1* mutant, suggesting a potential dependence of HopF2*_Pto_*-HA on QSK1 for both its accumulation and functionality in plants. Furthermore, RT-qPCR analysis showed comparable DEX-induced expression of *HopF2_Pto_-HA* in both *pDEX:HopF2_Pto_-HA* and *qsk1/ pDEX:HopF2_Pto_-HA* lines (Fig.10B), suggesting that the dependency of HopF2*_Pto_* on QSK1 is likely at the protein level rather than transcriptionally.

**Figure 10.**
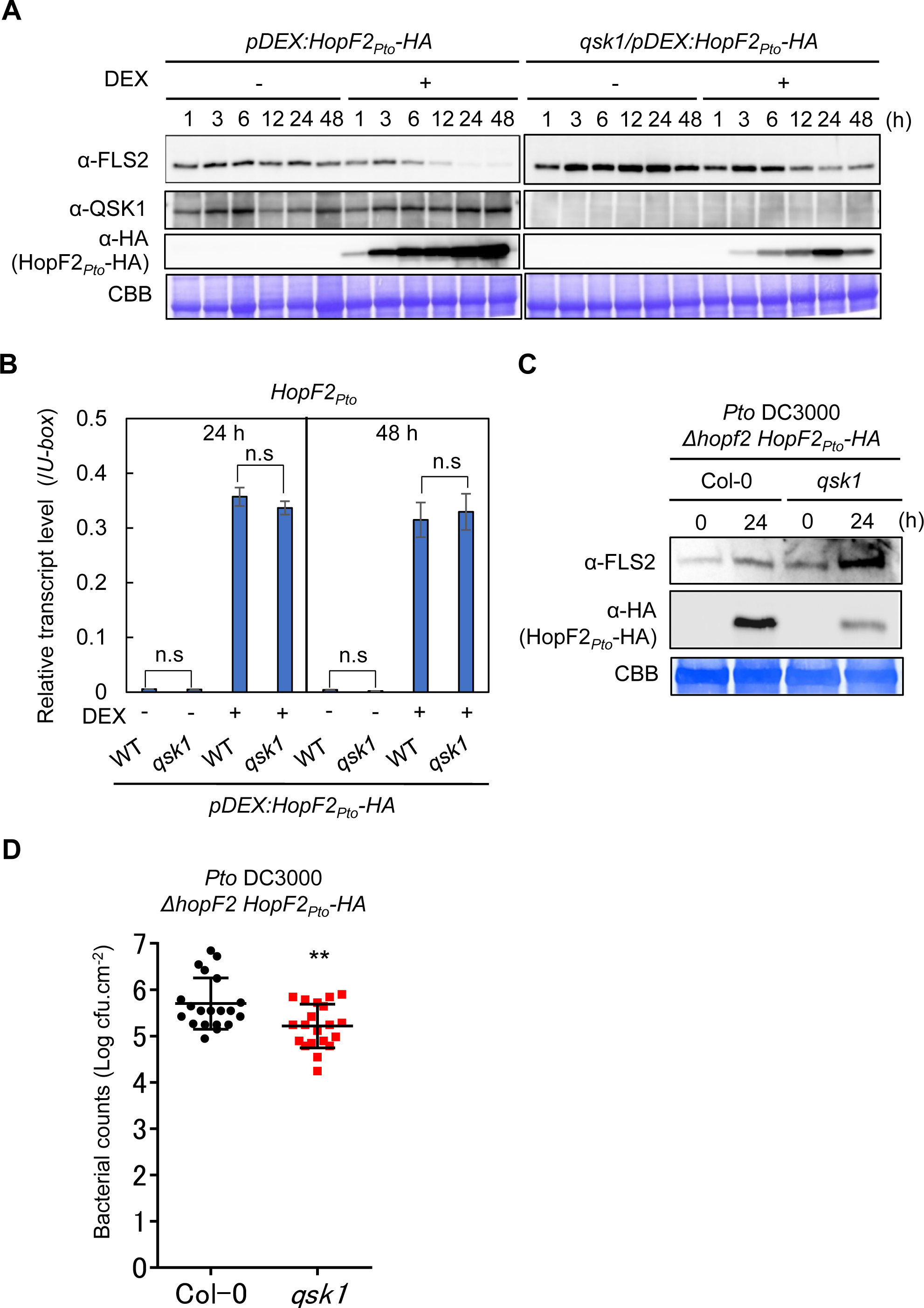
HopF2*_Pto_* requires QSK1 for its protein accumulation and function. **A)** HopF2*_Pto_* requires QSK1 for its protein accumulation and suppression of FLS2 accumulation. Two-week-old Arabidopsis seedlings of *pDEX:HopF2_Pto_-HA* and qsk1/*pDEX:HopF2_Pto_-HA* were treated with 30 µM DEX for 1, 3, 6, 12, 24 and 48 h and FLS2, QSK1, and HopF2*_Pto_*-HA protein levels were measured by immunoblotting with α-FLS2, α-QSK1, and α-HA antibodies. **B)** QSK1 does not affect *HopF2_Pto_* transcript levels. Transcript levels of *HopF2_Pto_-HA* in two-week-old Arabidopsis seedlings of *pDEX:HopF2_Pto_-HA* and *qsk1*/*pDEX:HopF2_Pto_-HA* treated with 30 µM DEX for 24 h and 48 h were measured by RT-qPCR after normalization to the *U*-*box* housekeeping gene transcript (*At5g15400*). erisks indicate significant differences (Student’s t-test, \*\**p* ≤ 0.01). All the experiments were repValues are mean ± SE of three biological replicates. There are no significant differences at *p* ≤ 0.05 between the two lines with or without treatment with DEX (Student’s t-test). **C)** HopF2*_Pto_* requires QSK1 during infection. Five-week-old Arabidopsis Col-0 and *qsk1* mutant were syringe-inoculated with *Pto* DC3000 *Δhopf2 HopF2_Pto_-HA* (inoculum: 10^8^ cfu/ml). Immunoblotting detecting FLS2 and HopF2*_Pto_*-HA at 1 d post-infection (dpi). The similar bacterial population at 1 dpi was confirmed by the bacterial growth assay shown in Supplemental Figure S13. **D)** *qsk1* mutant is more resistant against *Pto* DC3000 *Δhopf2 HopF2_Pto_-HA*. *Pto* DC3000 *Δhopf2 HopF2_Pto_-HA* were sprayed onto leaf surfaces of five-week-old soil-grown Arabidopsis plants at a concentration of 1x10^5^ cfu/mL. Data are means ± standard deviation of 20 replicates. The central horizontal line indicates the mean value. Asteated three times with similar results.

To understand this functional relationship during infection, we introduced HopF2*_Pto_*-HA into the *Pto* DC3000 strain and subsequently infected both Col-0 and *qsk1* mutants. At 24 h post-inoculation, HopF2*_Pto_*-HA accumulated more in Col-0 than in the *qsk1* mutant (Fig. 10C), while FLS2 levels were lower in Col-0 relative to the *qsk1* mutant when infected with *Pto* DC3000 harboring *HopF2_Pto_-HA*. Importantly, bacterial populations remained consistent between Col-0 and *qsk1* mutants at this time point (Supplemental Fig. S13). These findings strengthen our hypothesis that QSK1 is necessary for maintaining HopF2*_Pto_*’s protein stability and its ability to diminish FLS2 protein during infection. The *qsk1* mutant was more resistant against *Pto* DC3000 *ΔhopF2 HopF2_Pto_-HA* than Col-0 at 3 dpi, further supporting the importance of QSK1 in stabilizing and facilitating HopF2*_Pto_*’s function during infection (Fig. 10D).

## Discussion

In this study, we addressed the critical need for plants to precisely control the activity of PRR complexes, a safeguard against the detrimental outcomes of unexpected or excessive immune activation. We discovered QSK1 as a novel modulator of these complexes, primarily through its influence on the abundance of PRR proteins. Notably, our findings reveal an interaction between the type-III effector HopF2*_Pto_* and QSK1, which is pivotal for the stabilization of HopF2*_Pto_* within plants. Once stabilized by QSK1, HopF2*_Pto_* effectively inhibits SCOOP phytocytokine signaling and downregulates the cell’s responses to PAMPs and DAMPs by reducing PRR protein levels (Supplemental Fig. S14).

### QSK1 negatively regulates PTI through modulation of PRR protein levels

A tomato homolog of QSK1, TOMATO ATYPICAL RECEPTOR-LIKE KINASE 1 (TARK1), acts as a negative regulator of immunity as shown by increased resistance to pathogens in *tark1*-knockout lines and enhances susceptibility in its overexpression lines (Guzman et al., 2020). This indicates a conserved role of QSK1 in PTI across species. In Arabidopsis, QSK1-like proteins, LRR1, RKL1, and RLK90 may similarly modulate PTI (Supplemental Fig. S15), supported by elevated ROS production in *lrr1* and *rkl1* mutants in response to flg22, elf18, and pep1, a phenotype shared with the *qsk1* mutant. These results suggest that these homologs may function redundantly with QSK1 in PTI.

QSK1, also known as ALK1 (Auxin-induced LRR Kinase1) and KIN7 (Kinase 7), influences channels and transporters through phosphorylation, such as activating the TPK1 potassium channel during stomatal closure (Isner et al., 2018), and modifying the ABC transporter ABCG36, which affects export of the auxin precursor indole-3-butyric acid and the phytoalexin camalexin (Aryal et al., 2023). QSK1 is also involved in drought stress responses (Chen et al., 2021) and the regulation of callose-mediated plasmodesmata regulation and lateral root development during osmotic stress (Grison et al., 2019). Our study demonstrated an additional role for QSK1 in PTI regulation, by modulating PRR abundance, distinct from its known pathways. QSK1 was also shown to function as a co-receptor of Sucrose-induced Receptor Kinase 1 (SIRK1), facilitating the phosphorylation and activation of aquaporin PIP2;4 upon recognition of endogenous pep7 peptides (Wu et al., 2019; Wang et al., 2022). Our experiments showed no significant impact of this pathway on PTI responses (Supplemental Fig. S16), suggesting that FLS2 modulation by QSK1 does not depend on the pep7-SIRK1 signaling pathway.

We observed that *QSK1* overexpression leads to a reduction of FLS2 protein levels at the plasma membrane (Fig. 5). Additionally, ConA inhibits the QSK1-mediated reduction of both EFR and FLS2, implying that QSK1 induces the vacuolar degradation of the PRRs through autophagy or endocytosis (Fig. 5G). This aligns with recent findings showing that the LRR-RK ROOT MERISTEM GROWTH FACTOR 1 INSENSITIVE (RGI) recognizes the phytocytokine peptide GOLVEN2 (GLV2) and interacts with FLS2, enhancing its protein levels (Stegmann et al., 2022). Interestingly, the RGI3 ectodomain directly interacts with that of QSK1 and RLK902, and RGI4 ectodomain interacts with RKL1 *in vitro* (Smakowska-Luzan et al., 2018). The interaction between QSK1, RGIs, and their homologs might imply a complex interplay that disrupts the GLV2-mediated interaction between RGI and FLS2. Such disruption could cause the degradation of GLV2-unbound FLS2. A comprehensive understanding of the intricate relationship between phytocytokine signaling and FLS2 homeostasis, especially QSK1’s involvement remains a critical area for future research.

### HopF2*_Pto_* decreases plant responsiveness to PAMPs, DAMPs and SCOOP phytocytokines by reducing PRR levels

Previous studies have established HopF2*_Pto_*as a potent inhibitor of PTI responses such as ROS production, MAPK activation, and callose deposition (Wu et al., 2011; Hurley et al., 2014; Zhou et al., 2014). Our work shows an additional role for HopF2*_Pto_* in diminishing plant response to PAMPs, DAMPs, and SCOOP phytocytokines specifically through reducing PRR levels and *PROSCOOPs* transcript levels. Interestingly, HopU1, another effector encoding a MARylation enzyme from *Pto* DC3000 also modulates FLS2 protein levels, by targeting GRP7, an RNA-binding protein in FLS2 translation (Fu et al., 2007; Nicaise et al., 2013). Unlike HopU1, which does not affect steady-state FLS2 levels (Nicaise et al., 2013), HopF2*_Pto_* significantly reduces both baseline (Fig. 6, C to D) and post-infection FLS2 levels (Fig. 9B). Thus, *Pto* DC3000 employs these two distinct MARlyation enzyme-coding effectors to manipulate FLS2 regulation in various ways. Furthermore, the pathogen uses the ubiquitin ligase AvrPtoB to degrade FLS2 by polyubiquitinating its kinase domain (Goehre et al., 2008). These strategies collectively highlight the significance of PRR suppression in the virulence mechanism of pathogens like *Pto* DC3000.

### The Interplay of HopF2*_Pto_* with MIK2 and PRR expression in modulating plant immunity responses

HopF2*_Pto_* significantly reduces the transcript levels of important PRRs, including FLS2, LORE, CARD1, RDA2, and MIK2, as well as a majority of *PROSCOOPs*. Intriguingly, *mik2* mutants exhibit reduced flg22-triggered ROS production (Rhodes et al., 2021), hinting at MIK2’s role in maintaining baseline expression of *FLS2* and *PROSCOOPs*, through subtle activation by SCOOP peptides. This is further supported by the findings that MIK2 activation by SCOOP12 increases *FLS2* and PROSCOOP transcripts (Hou et al., 2021). Therefore, HopF2*_Pto_*’s impact on FLS2 levels might involve disrupting this MIK2-dependent positive feedback loop. However, the HopF2*_Pto_*-induced reduction in FLS2 cannot be solely attributed to MIK2 disruption. This is evident as HopF2*_Pto_* expression completely inhibits flg22-induced responses, whereas *mik2* mutants still retain some responsiveness to flg22 (Supplemental Figs. S11 and S12)(Rhodes et al., 2021).

### Distinct mechanisms of PRR degradation by HopF2*_Pto_* : Exploring vacuolar degradation and transcript regulation

ConA’s inhibition of the *HopF2_Pto_*-induced EFR reduction implies that HopF2*_Pto_* might target EFR for vacuolar degradation. However, ConA does not reverse HopF2*_Pto_*’s reduction of FLS2 protein (Supplemental Fig. S8A), possibly attributed to HopF2*_Pto_*’s differential effects on their transcripts: steady-state FLS2 transcripts are diminished, while EFR transcripts remain unaffected. Consequently, even if ConA inhibits the vacuolar degradation of FLS2, the diminished levels of FLS2 transcripts may still limit its protein synthesis. In contrast, EFR protein loss under HopF2*_Pto_*might be mainly through vacuolar degradation. This distinction is highlighted by the more pronounced reduction of FLS2 and FLS2-mediated MAPKs activation than EFR by HopF2*_Pto_* (Figs. 6 and 9; Supplemental Fig. S12).

Previous studies have shown that signaling-inactive FLS2 undergoes degradation through selective autophagy with Orosomucoid (ORM) proteins as key autophagy receptors (Yang et al., 2019), while signaling-active FLS2 undergoes vacuolar degradation through endocytosis (Robatzek et al., 2006; Beck et al., 2012; Mbengue et al., 2016). HopF2*_Pto_*might exploit either pathway to diminish PRR protein levels. Despite our hypotheses, direct observation of EFR or FLS2 within autophagosomes or endosomes after expressing *HopF2_Pto_* was not feasible, likely due to the low expression levels of *EFR-GFP* in our Arabidopsis transgenic lines (*pEFR:EFR-GFP*) and the reduced FLS2 transcript levels complicating detailed microscopic observation of FLS2-GFP in *pFLS2:FLS2-GFP* lines.

### The MARylation activity of HopF2*_Pto_* is required for its virulence

Our finding establishes the critical role of HopF2*_Pto_*’s catalytic residue in MARylation for FLS2 protein reduction (Fig. 7). However, the exact mechanisms through which HopF2*_Pto_*influences transcriptome reprogramming changes and vacuolar degradation of PRRs via MARylation remain elusive. Previous studies demonstrated that HopF2*_Pto_* targets key regulators of the PTI signaling pathway, including MKK5 and BAK1 (Wang et al., 2010; Zhou et al., 2014), as well as RIN4 (Wilton *et al*., 2010), impacting both PTI and ETI. It is plausible that HopF2*_Pto_*-mediated inhibition of MKK5 and BAK1 contributes to transcriptome reprogramming, possibly by disrupting MIK2 activation by SCOOP peptides (Hou et al., 2021; Rhodes et al., 2021). However, MKK5 and BAK1 are unlikely candidates for HopF2*_Pto_*-mediated autophagy and/or endocytosis of PRRs, because both proteins are not part of a stable PRR complex in the absence of PAMP treatment (Chinchilla et al., 2007). Instead, HopF2*_Pto_*may MARylate other proteins to induce autophagy and/or endocytosis of PRRs.

We propose several hypotheses for HopF2*_Pto_*induction of PRR degradation. Firstly, HopF2*_Pto_* may MARylates and activates QSK1. This activation could inhibit the RGI-FLS2 association, leading to PRR destabilization and their subsequent degradation through autophagy and/or endocytosis. This hypothesis is supported by the fact that both QSK1 and HopF2*_Pto_* induce vacuolar degradation of PRRs (Figs 5, 6, and 9). Another hypothesis is that HopF2*_Pto_* directly MARylates PRRs, altering their structural conformation to enhance ORM protein binding and thus autophagy. In this scenario, QSK1 might serve as a scaffold, facilitating PRR MARylation. Lastly, HopF2*_Pto_*might target G proteins, known to inhibit FLS2 autophagy (Miller et al., 2019). This is supported by the fact that bacterial toxins predominantly MARylate Gα proteins in animals (Ishiwata-Endo et al., 2020). Detecting HopF2*_Pto_*’s MARylation *in vivo* remains technically challenging, particularly direct observation of the MARylation of QSK1 and PRRs. Future studies should focus on identifying proteins MARylated by HopF2*_Pto_ in vivo* and clarifying their roles in the vacuolar degradation of PRRs through autophagy and/or endocytosis.

### HopF2*_Pto_* requires QSK1 for its stabilization and function

Our findings indicate that QSK1 plays a pivotal role in stabilizing HopF2*_Pto_* in plants, although its exact mechanism remains elusive. Notably, HopF2*_Pto_*, known to possess a predicted myristoylation sequence essential for plasma membrane localization and virulence (Wilton et al., 2010), seems to stabilize when it interacts with QSK1, following myristoylation. This interaction may assist HopF2*_Pto_*in targeting the PRR complex. The complex interplay between QSK1 and HopF2*_Pto_*, while not fully understood, indicates a broader role for QSK1 and its homologs in aiding virulence effectors across various plants. For instance, XopN, a virulence factor from *X. campestris*, interacts with TARK1, a tomato homolog of QSK1 (Kim et al., 2009; Guzman et al., 2020). In *tark1*-silenced plants, XopN’s virulence function is notably reduced, suggesting that TARK1 is crucial for XopN functionality. Moreover, TARK1 may guide XopN to interact with tomato 14-3-3 isoform TFT1, a positive regulator of PTI in tomatoes (Taylor et al., 2012). This relationship mirrors that of HopF2*_Pto_*-QSK1-PRR, though it remains unclear if TARK1 primarily maintains XopN protein stability, and facilitates its integration into the PRR complex.

## Methods

### Plant materials and growth conditions

*Arabidopsis thaliana* (L.) Heynh. Plants were grown on soil under an 8 h or 16 h photoperiod at 23°C, or in a half-strength MS medium containing 1% sucrose under a continuous light photoperiod at 23°C. *Nicotiana benthamiana* Domin. Plants were soil-grown under a 16 h photoperiod at 25°C.

### Vector construction and generation

To generate epiGreenB5-*p35S:QSK1-3×HA*, and epiGreenB5-*p35S:QSK1-GFP*, CDS region of QSK1 was amplified by PCR with KoD FX neo (Toyobo, Osaka, Japan) and the resulting PCR product was cloned into the epiGreenB5 (3*x*HA) and epiGreenB (eGFP) vectors between the *Cla*I and *Bam*HI restriction sites with an In-Fusion HD Cloning Kit (Clontech, CA, USA) (Nekrasov et al., 2009). To generate epiGreenB5-*pQSK1:QSK1-GFP,* an amplicon containing the 2000-bp promoter upstream of the start codon and the coding regions of QSK1 was cloned into the epiGreenB (eGFP) vectors between the *EcoRI* and *Bam*HI restriction sites with In-Fusion HD Cloning Kit. pCAMBIA2300-*pFLS2:FLS2-GFP* was described previously (Robatzek et al., 2006).

### Transgenic lines and T-DNA insertion lines

Arabidopsis stable transgenic lines of *p35S:QSK1-3×HA* (epiGreenB5), *p35S:QSK1-GFP* (epiGreenB5), and *qsk1/pQSK1:QSK1-GFP* (epiGreenB5) were generated by the floral drop and floral dip methods. T-DNA insertion mutant lines, *qsk1* (SALK_ 019840C), *lrr1* (WiscDsLoxHs082_03E), *rkl1* (SALK_099094C), *sirk1* (SALK_125543C), and *pep7* (SALK_025824C) were obtained from the Arabidopsis Biological Resource Center at the Ohio State University. Previously published lines were: *bak1bkk1* (Roux et al., 2011), *fls2*, *pFLS2:FLS2-GFP* (Robatzek et al., 2006), *efr-1/pEFR:EFR-GFP*, *rbohD/pRBOHD:3xFLAG-gRBOHD* (Kadota et al., 2014), *pDEX:HopF2_Pto_-HA* and its variant D175A (Wilton et al., 2010). Homozygous *pFLS2:FLS2-GFP/p35S:QSK1-3xHA, pEFR:EFR-GFP/p35S:QSK1-3xHA, qsk1*/*pDEX:HopF2_Pto_*-HA, *pDEX:HopF2_Pto_-HA*/*/pFLS2:FLS2-GFP,* and *pDEX:HopF2_Pto_*-HA/*/pEFR:EFR-GFP* lines were generated by crossing homozygous lines and then selection by genotyping.

### Generation of QSK1 antibody

A polyclonal anti-QSK1 antibody was produced by immunizing rabbits with a synthetic peptide (NH2-C+EEVSHSSGSPNPVSD-COOH) originating from the C-terminal region of QSK1 (Eurofins Scientific SE, Luxembourg).

### Immunoblotting

Immunoblotting was performed with antibodies diluted in the blocking solution (5% nonfat milk in TBS with 0.1% [v/v] Tween) at the following dilutions: α-GFP antibody (ab290, Abcam, Cambridge, UK), 1:8,000; α-HA-horseradish peroxidase (HRP) (3F10, Roche, Basel, Switzerland), 1:5,000; α-FLAG-HRP (M2 monoclonal antibody, Sigma-Aldrich, St. Louis, MO, USA), 1:2000; α-FLS2 (Chinchilla et al., 2006), 1:1,000; α-BAK1 (Roux et al., 2011), 1:1000; α-QSK1,1:500, and α-rabbit-HRP conjugated antibody (NA934; GE Healthcare, Chicago, IL, USA), 1:10,000. For detection of RBOHD, α-RBOHD (AS152962; 1:1,000; Agrisera, Vännäs, Sweden) antibody was diluted in Can Get Signal® Solution 1 (Toyobo, Osaka, Japan) and the α-rabbit-HRP conjugated antibody was diluted in Can Get Signal® Solution 2 to enhance the signal of immunoblotting.

### Bacterial strains

*Pto* DC3000 *ΔhopF2 HopF2_Pto_-HA* was described previously (Wilton et al., 2010). It is important to note that the native HopF2*_Pto_* has an ATA start codon, which limits its expression. On the other hand, *Pto* DC3000 *ΔhopF2 HopF2_Pto_- HA* uses the more common ATG start codon, resulting in enhanced expression of *HopF2_pto_-HA* during the infection. To generate *Pseudomonas fluorescens* (Pf0-1) *HopF2_Pto_-HA* and *P. fluorescens* Pf0-1*HopF2_Pto_* (D175A)-*HA*, *P. fluorescens* Pf0-1 was transformed with the expression vectors, *schF2/hopF2_Pto_ ^ATG^:HA* or *schF2/hopF2_Pto_ ^ATG^* (D175A)*:HA*.

### Statistical Analysis

Statistical significances based on t-test and one-way ANOVA were determined with GraphPad Prism6 software (GraphPad Software, San Diego, CA, USA). Statistical data are provided in Supplemental Data Set S7.

### Other methods

Chemical inhibitors were described in Methods S1. Protein extraction, IP, protein identification by LC-MS/MS, ROS burst assay, MAPK activation assay, bacterial infection assays, phylogenetic analyses, transient expression in *N*. *benthamiana,* confocal microscopy analyses, RT-qPCR assay, QIS-Seq analyses, RNA-seq and differential gene expression analyses, PCA with SOM clustering, and GO term enrichment analyses were performed as described previously (Lewis et al., 2012; Kadota et al., 2014; Goto et al., 2020; Goto et al., 2023) with minor modifications detailed in Supplemental Methods S1. All primers used in this study are listed in Supplemental Data Set S8.

## Acknowledgments

We thank all members of the Shirasu lab for discussion. We thank Ayami Furuta, Naomi Watanabe, Mamiko Kouzai, Mizuki Yamamoto, and Yoko Nagai for their support of this project.

## Author contributions

YK and MM performed IP experiments. JS, PD, and FLHM. performed LC-MS/MS analyses. NM helped to generate plasmids. YK, YG, and HM characterized the phenotype of the *qsk1* mutant and overexpression lines. JDL performed QIS seq. YK and YG analyzed the effect of HopF2*_Pto_*on FLS2 homeostasis and the role of QSK1 for HopF2*_Pto_* function. AS, TS, YI performed RNA-seq analyses and YK and YG analyzed the data. YK, DSG, HN, SR, DD, CZ, and KS supervised the research. YK and YG wrote the draft manuscript. All the authors commented on the manuscript.

## Funding

The research was financially supported by JSPS KAKENHI Grant Numbers 16J00771 (to Y.G), 16H06186, 16KT0037, 20H02994, 21K19128 (to Y.K), 17H06172, 20H05909, 22H00364 (to K.S), USDA ARS 2030-21000-046-00D and 2030-21000-050-00D (JDL), as well as the Gatsby Charitable Foundation (to F.L.H.M, C.Z., and S.R.), the Natural Sciences and Engineering Research Council of Canada (DD and DSG), and the European Research Council (project ‘PHOSPHinnATE’, grant agreement No. 309858 to C.Z. and project “STORM”, grant agreement No. 311310 to S.R.).

## Data availability

The data underlying this article are available in the article and in its online supplementary material.

## Supporting Information

**Supplemental Figure S1.** Heterologous expression of *QSK1-3xHA* reduces flg22-induced ROS production in *Nicotiana benthamiana*.

**Supplemental Figure S2.** T-DNA insertion and expression in *qsk1* mutant.

**Supplemental Figure S3.** Phenotype recovery in *qsk1* complementation lines.

**Supplemental Figure S4.** *QSK1* overexpression lines are slightly smaller than Col-0 and *qsk1* mutant.

**Supplemental Figure S5.** QSK1 localizes at the plasma membrane.

**Supplemental Figure S6.** flg22 and elf18 induce the accumulation of *QSK1* transcript.

**Supplemental Figure S7.** Pharmacological analyses of FLS2 reduction induced by QSK1.

**Supplemental Figure S8.** Pharmacological analyses of FLS2 reduction induced by HopF2*_Pto_*.

**Supplemental Figure S9.** Multidimensional scaling (MDS) plot with self-organizing map (SOM) clustering of genes affected by HopF2*_Pto_*.

**Supplemental Figure S10.** HopF2*_Pto_* affects some transcript levels of commonly associated proteins with EFR, FLS2, and RBOHD.

**Supplemental Figure S11.** HopF2*_Pto_* inhibits PAMP-induced ROS production.

**Supplemental Figure S12.** HopF2*_Pto_* inhibits PAMP-induced activation of MAPKs.

**Supplemental Figure S13.** *Pto* DC3000 *Δhopf2 HopF2_Pto_-HA* grows to the same extent in Col-0 and *qsk1* mutant.

**Supplemental Figure S14.** A model of the virulence function of HopF2*_Pto_* suppressing PTI and its relationship to QSK1.

**Supplemental Figure S15.** *lrr1* and *rkl1* mutants have higher ROS production in response to flg22, elf18, and pep1.

**Supplemental Figure S16.** SIRK1 and PEP7 do not affect PAMP-induced ROS production.

**Supplemental Methods S1.** Additional methods.

**Supplemental Data Set S1_1.** FLS2-associated proteins.

**Supplemental Data Set S1_2.** EFR-associated proteins.

**Supplemental Data Set S1_3.** RBOHD-associated proteins.

**Supplemental Data Set S1_4.** Commonly associated proteins with FLS2, EFR, and RBOHD.

**Supplemental Data Set S2_1.** Peptide counts of QSK1 in FLS2-GFP IP analysis.

**Supplemental Data Set S2_2.** Peptide counts of QSK1 in EFR-GFP IP analysis.

**Supplemental Data Set S2_3.** Peptide counts of QSK1 in FLAG-RBOHD IP analysis.

**Supplemental Data Set S3_1.** Enrichment score of HopF2*_Pto_* interactors in Quantitative Interactor Screening with Next-Generation Sequencing (QIS-Seq).

**Supplemental Data Set S3_2.** In planta proximity-dependent biotin identification (BioID) of HopF2*_Pto_* associated proteins

**Supplemental Data Set S4_1.** Normalized expression values of genes in Arabidopsis seedlings of Col-0 or *pDEX:HopF2_Pto_-HA* lines treated with DMSO or DEX.

**Supplemental Data Set S4_2.** Up-regulated genes by DEX treatment compared to DMSO treatment in Col-0 (log2 fold change ≥ 1, FDR ≤ 0.05).

**Supplemental Data Set S4_3.** Down-regulated genes by DEX treatment compared to DMSO treatment in Col-0 (log2 fold change ≤ -1, FDR ≤ 0.05).

**Supplemental Data Set S4_4.** Up-regulated genes by the expression of *HopF2_Pto_* (*pDEX:HopF2_Pto_+DEX* vs Col-0+DEX) (log2 fold change ≥ 1, FDR ≤ 0.05).

**Supplemental Data Set S4_5.** Down-regulated genes by the expression of *HopF2_Pto_* (*pDEX:HopF2_Pto_*+DEX vs Col-0+DEX) (log2 fold change ≤ -1, FDR ≤ 0.05).

**Supplemental Data Set S4_6.** Up-regulated genes by DEX treatment in *pDEX:HopF2_Pto_-HA* line (*pDEX:HopF2_Pto_*+DEX vs *pDEX:HopF2_Pto_*+DMSO) (log2 fold change ≥ 1, FDR ≤ 0.05).

**Supplemental Data Set S4_7.** Down-regulated genes by DEX treatment in *pDEX:HopF2_Pto_-HA* line (*pDEX:HopF2_Pto_*+DEX vs *pDEX:HopF2_Pto_*+DMSO) (log2 fold change ≤ -1, FDR ≤ 0.05).

**Supplemental Data Set S5_1.** Gene ontology (GO) enrichment analysis of the up-regulated genes by HopF2*_Pto_*

**Supplemental Data Set S5_2.** Gene ontology (GO) enrichment analysis of the down-regulated genes by HopF2*_Pto_*

**Supplemental Data Set S5_3.** Gene ontology (GO) enrichment analysis of the genes in the cluster 1

**Supplemental Data Set S5_4.** Gene ontology (GO) enrichment analysis of the genes in the cluster 2

**Supplemental Data Set S6.** Genes in SOM clusters.

**Supplemental Data Set S7.** Results of statistical analysis.

**Supplemental Data Set S8.** Primers used in this paper.

